# Adaptation to transient and local electrical muscle stimulation elicits persistent and global changes in walking kinematics

**DOI:** 10.1101/2025.11.05.686889

**Authors:** Ryosuke Murai, Daichi Nozaki

## Abstract

It is generally assumed that motor commands altered by adaptation to a novel environment revert to their original state once the environment returns to baseline. However, this assumption—based mainly on simple movements such as reaching—may not hold for complex actions. In redundant systems, de-adaptation can cause motor commands and resulting kinematics to settle into states distinct from the original. Here, we examined adaptation and de-adaptation in treadmill walking while modulating lower-limb dynamics using closed-loop neuromuscular electrical stimulation (NMES) applied bilaterally to the tibialis anterior. Adaptation to a localized ankle perturbation induced global kinematic changes across the ankle, knee, and hip joints. After perturbation removal, joint kinematics did not fully return to baseline; instead, features of the adapted gait persisted for eight minutes. These findings highlight the distinctive nature of motor adaptation in redundant systems and demonstrate the potential of NMES-based perturbations for inducing implicit modifications in movement patterns.

## Introduction

To consistently perform accurate movements, the motor system automatically and unconsciously adjusts motor commands by integrating error information from multiple sensory modalities [Wolpert et al., 1995; Shadmehr et al., 2010]. This process, known as implicit motor adaptation, ensures precise motor execution amid ever-changing external environments and bodily states.

The mechanisms underlying this process have been extensively studied in visually guided reaching tasks, where behavioral and neurophysiological changes have been examined in response to external perturbations. In a typical reaching experiment, participants perform hand-reaching movements toward a target while their movements are disturbed by mechanical or visual perturbations [Shadmehr and Mussa-Ivaldi, 1994; Thoroughman and Shadmehr, 2000; Cunningham, 1989; Krakauer, 2009]. These perturbations initially cause a sudden increase in movement errors (e.g., deviations from the target direction in visual space); however, these errors gradually decrease as the sensorimotor system adapts to the novel condition. When the perturbation is removed after adaptation occurs (often termed the washout phase or post-adaptation phase), movement errors appear in the opposite direction to that observed during the imposition of the perturbation. These errors, referred to as aftereffects, are considered evidence of implicit adaptation [Shadmehr and Mussa-Ivaldi, 1994; Mazzoni and Krakauer, 2006; Krakauer, 2009; Taylor et al., 2014].

In standard reaching tasks, aftereffects typically diminish monotonically over time, with movement kinematics gradually returning to baseline as the acquired adaptation decays. This behavior likely reflects the kinematically non-redundant nature of such tasks: the reaching movements examined in these experiments are generally planar, involving primarily the shoulder and elbow joints. Consequently, the degrees of freedom in the task space (two-dimensional hand position) correspond directly to those in the control space (shoulder and elbow joint angles). This lack of redundancy ensures that the joint configuration required to achieve the reaching movement remains essentially identical before and after the perturbation, at least at the level of joint kinematics [Kobayashi and Nozaki, 2024].

While reaching movements in laboratory settings are typically kinematically non-redundant, many real-world motor behaviors, such as walking, are inherently redundant. This raises an important question: do joint kinematics in such redundant movements return to baseline after the removal of perturbations, as observed in reaching tasks? Because the musculoskeletal system allows an infinite number of joint configurations to achieve the same goal, it is theoretically possible that kinematics differ before and after exposure to a perturbation.

Addressing this question offers insight into a fundamental problem in motor control—how the nervous system selects a specific movement pattern from countless possibilities to achieve a desired outcome [Bernstein, 1996]. Previous studies have proposed that the motor system selects commands that optimize movement according to certain cost functions [Flash and Hogan, 1985; Harris and Wolpert, 1998; Todorov and Jordan, 2002; Todorov, 2004; Izawa et al., 2008; Emken et al., 2007; Finley et al., 2013]. Within this framework, if the system consistently identifies a global optimum, kinematics during washout should match baseline even in redundant tasks. Conversely, if multiple suboptimal solutions exist near the optimum, the resulting kinematics after perturbation removal may deviate from the original state. Thus, examining joint kinematics during washout in redundant movements provides a unique window into how the motor system resolves the ill-posed problem imposed by musculoskeletal redundancy.

Despite this potential importance, joint kinematics during the washout phase have not been a primary focus of much of the prior research on redundant motor tasks. In gait adaptation, for example, a large proportion of studies have utilized the split-belt treadmill adaptation paradigm, where participants adapt to a belt speed difference between legs [Dietz et al., 1994; Choi and Bastian, 2007; Reisman et al., 2005; Finley et al., 2013]. In this paradigm, much prior research has focused on summary metrics (e.g., step length asymmetry) as indicators of adaptation, rather than examining individual joint motion. While these summary parameters generally return to baseline level during the washout phase, similar to movement errors in reaching tasks, they do not directly capture possible changes in kinematics. In addition to split-belt treadmills, mechanical perturbations using robots or exoskeletons have also been used to study gait adaptation [Selinger et al., 2015; Cajigas et al., 2017; Emken et al., 2007; Gordon and Ferris, 2007]. For example, Selinger et al. investigated adaptive changes in step frequency in response to a mechanical perturbation that shifted the energetically optimal step frequency away from baseline [Selinger et al., 2015]. They found that even after the perturbation was removed, the step frequency remained close to that established under the perturbation for several minutes, even though it was no longer energetically optimal. Again, however, step frequency is a summary measure and does not reflect the redundancy of the musculoskeletal system. Additionally, although some studies using mechanical perturbations have investigated changes in joint kinematics, the focus was generally placed on adaptation-induced changes rather than on their behavior during the washout phase [Gordon and Ferris, 2007; Gordon et al., 2013].

In this study, we aimed to investigate changes in gait kinematics induced by adaptation to a movement perturbation, with a particular focus on the washout process. To achieve this, we employed a novel perturbation system using closed-loop neuromuscular electrical stimulation (NMES; hereinafter, electrical stimulation above the motor threshold is referred to as NMES), which we recently proposed as a method for systematically perturbing whole-body movements with minimal interference to voluntary motion [Murai et al., in preparation]. In this system, NMES is systematically applied with an intensity modulated by real-time movement patterns to induce involuntary muscle contractions, thereby altering the relationship between motor commands and limb movements. This closed-loop NMES effectively alters musculoskeletal dynamics, providing a controlled means to study motor adaptation in whole-body movements. Using this perturbation system, we manipulated the musculoskeletal dynamics of the lower limbs during treadmill walking by applying NMES bilaterally to the tibialis anterior (TA). The intensity of NMES was modulated based on the ankle plantarflexion angle during walking. This approach enabled us to examine how the sensorimotor system adjusts the original gait pattern both during perturbation exposure and in the subsequent washout phase.

## Methods

### Participants

A total of 16 males participated in this study (27.1 ± 5.2 years, 170.2 ± 5.9 cm, 70.0 ± 13.8 kg). The participants had no musculoskeletal or neurological disorders that could impair normal gait. Each participant provided written informed consent before participating in the experiments. The Ethics Committee of the University of Tokyo reviewed and approved the experimental protocol, which was in accordance with the Declaration of Helsinki.

### Experimental protocol

First, the participants practiced walking on a split-belt instrumented treadmill (Bertec Corporation, USA) until they felt accustomed to it (familiarization session). During this session, we measured the ankle dorsiflexion/plantarflexion over the gait cycle to determine the individual-specific function used to modulate NMES intensity (see *Perturbation*). After the familiarization session, participants performed a 19-minute walking task on the treadmill, which consisted of three phases: baseline (3 minutes), adaptation (8 minutes), and washout (8 minutes) (Fig. 1a). NMES perturbation was applied during the adaptation phase. Within the 8-minute adaptation phase, four 10-second catch blocks were interleaved at 1, 3, 5, and 7 minutes, during which the perturbation was temporarily switched off.

**Fig. 1.**
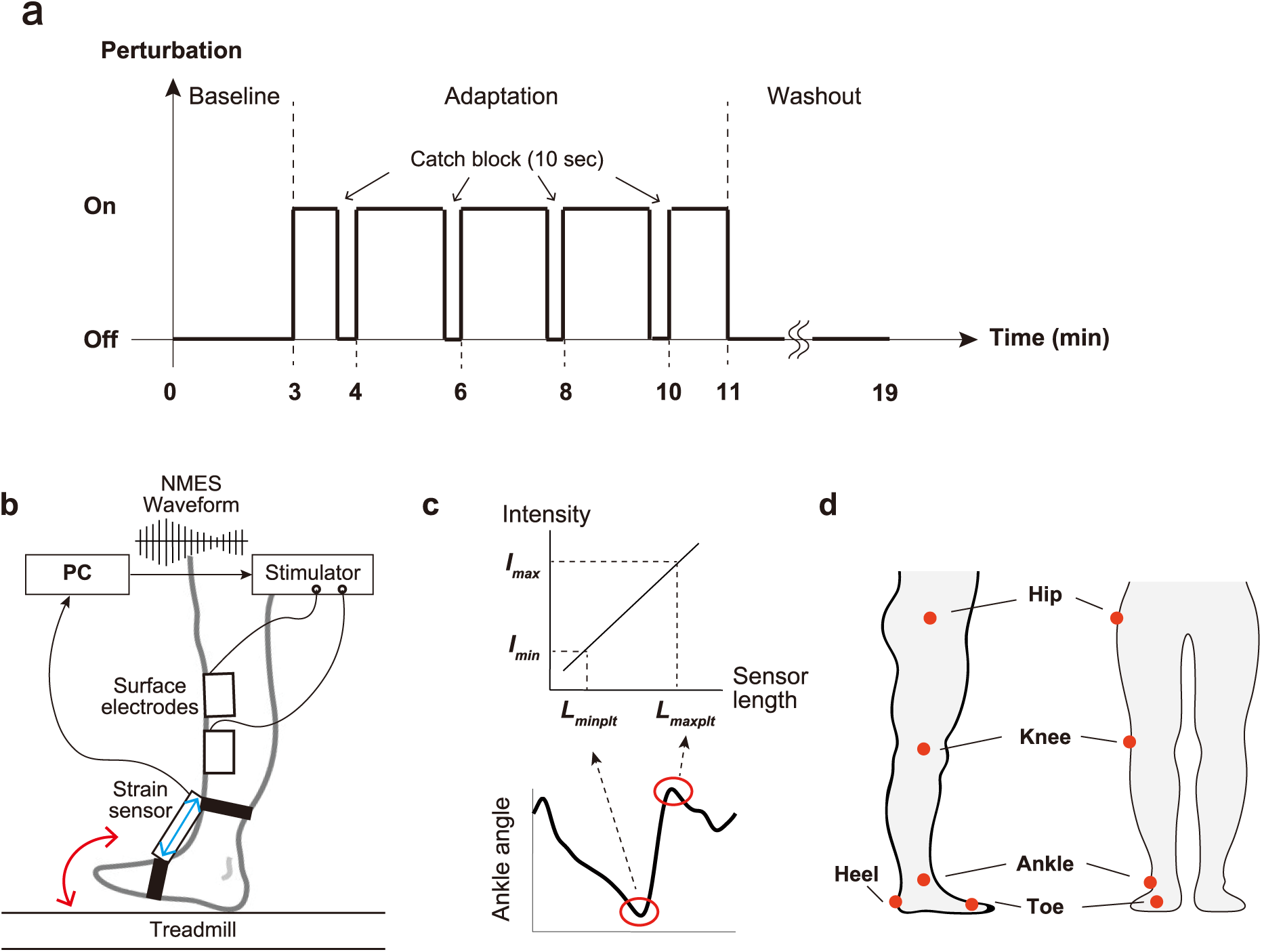
Experimental setup. (a) Perturbation schedule of the task. Participants walked on a treadmill wearing surface electrodes for electrical stimulation on both tibialis anterior muscles. (b) Closed-loop NMES perturbation system with an elastic strain sensor. The strain sensor was attached to the dorsum of foot and shank with hook-and-loop fasteners. Ankle dorsiflexion/plantarflexion (red arrow) was quantified by changes in the sensor length (blue arrow). The amplitude of the stimulation was modulated in real time based on the sensor length. (c) Individualized relationship between sensor length and stimulation intensity. The stimulation intensity was linearly modulated as a function of the sensor length. The slope and intercept of this relationship were determined for each participant, based on their sensitivity and tolerance to electrical stimulation (*I_max_* and *I_min_*) and the minimum and maximum sensor lengths in gait cycles measured during the familiarization session (*L_maxplt_*, and *L_minplt_*). (d) Marker placement used for joint angle calculations. Markers were placed bilaterally (only the markers on the right leg are shown in the figure).

The treadmill belt speed was maintained at 1.0 m/s throughout the experiment, and participants were not given specific instructions about their step frequency. They were instructed to walk naturally, without employing any explicit strategy or deliberately modifying their walking pattern in response to the perturbation.

### Perturbation

The participants’ gait was perturbed by closed-loop NMES applied to TA of both legs. The NMES intensity was modulated based on the ankle dorsiflexion/plantarflexion during gait, measured by an elastic strain sensor (C-STRETCH_®_, Bando Chemical Industries, Ltd., Japan) attached to the participants’ ankles (Fig. 1b). The sensor detected changes in the distance between its attachment points (the dorsum of the foot and the shank), and these changes were approximated as changes in joint angle.

The specific relationship between the ankle dorsiflexion/plantarflexion and stimulation intensity is illustrated in Fig. 1c. In brief, greater plantarflexion resulted in stronger stimulation. The NMES intensity was linearly manipulated as a function of the length of the strain sensor as follows:

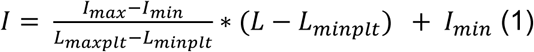

where *L* and *I* represent the sensor length and resultant stimulation intensity (i.e., the amplitude of the NMES pulse), respectively. *I_max_*, *I_min_*, *L_maxplt_*, and *L_minplt_* are individualized parameters for each participant. *I_max_* and *L_min_* corresponded to 60% of the highest tolerable intensity and the highest intensity that did not induce visible contraction of the TA, respectively. These values were determined for each leg in 2.5 mA increments before the task. *L_minplt_* and *L_minplt_* were the maximum and minimum sensor lengths within a stride, measured during the familiarization phase. To determine *L_minplt_* and *L_minplt_*, we extracted data from 15 consecutive strides after the participants became accustomed to treadmill walking during the familiarization session, and then calculated the median of the maximum and minimum values across these strides.

TA was stimulated using a linear isolator (ULI-100, Unique Medical Co., Ltd., Japan) through 5 × 9 cm surface electrodes (PALS_®_, Axelgaard Manufacturing Co., Ltd., USA). The electrodes were placed bilaterally at the proximal and distal ends of the TA muscle belly. The stimulation trains consisted of 500-μs biphasic, rectangular pulses, and were delivered at a frequency of 20 Hz, which is considered to induce less muscle fatigue than higher frequencies [Moritani et al., 1985].

### Data collection and analysis

Kinematic data were measured at 100 Hz using a Qualisys motion capture system (Qualisys AB, Sweden). The placement of reflective markers for joint angle calculations is shown in Fig. 1d. The hip, knee, and ankle joint angles were calculated as follows: the hip angle was defined as the angle between the thigh (the vector connecting the hip and knee markers) and the coronal plane; the knee angle was defined as the angle between the thigh and the shank (the vector connecting the knee and ankle markers); the ankle angle was defined as the angle between the shank and the foot (the vector connecting the toe and heel markers). Ground reaction force was collected at 1000 Hz by force sensors embedded in the treadmill to detect heel-strike and toe-off. Moments when the vertical force exceeded 50 N and when it dropped below 50 N were identified as heel-strike and toe-off, respectively. The interval from right heel-strike to the next right heel-strike was defined as a single stride. The marker and ground reaction force data were filtered using a second-order, zero-lag Butterworth filter with cutoff frequencies of 10 Hz and 30 Hz, respectively.

Out of the 16 participants, five were excluded due to incomplete data acquisition or errors in stimulation delivery. Additionally, one participant was excluded from the analysis due to low tolerance for NMES, which prevented sufficient muscle contraction. As a result, data from 10 participants were included in the analysis.

### Data analysis

All analyses were conducted using kinematic data from the right leg. The 19-minute task was segmented into 10-second blocks, and for each block, the joint kinematic data were averaged across strides that were fully contained within that block. Prior to averaging, the kinematic data for each stride were time-normalized. Baseline, late adaptation, and late washout gait patterns were defined as those of the last block of the baseline, adaptation, and washout phases, respectively. Additionally, to examine changes in joint kinematics at specific phases of the gait cycle, the joint kinematics within each block were divided into 10 bins, and the joint angles were averaged within each bin. These 10 bins corresponded to 0–10%, 10–20%, …, and 90– 100% of each gait cycle.

Simply averaging kinematic data across participants could obscure meaningful adaptive changes induced by the perturbation, due to differences in individual response patterns (see Results). To investigate changes in gait patterns during the task independently of inter-individual variability, we evaluated gait dissimilarity, defined as the extent to which gait patterns from different blocks differed within each participant. Specifically, we compared gait patterns in the washout phase with those of baseline and late adaptation to determine whether gait patterns returned to baseline following the removal of the perturbation. We quantified gait dissimilarity using the dynamic time warping (DTW) algorithm [Sakoe and Chiba, 1978] (Fig. 3a). Briefly, DTW is a method for measuring the similarity (dissimilarity) between two time series by optimally aligning them. The dissimilarity is calculated as the sum of the distances between the matched points in the two series, where matching is performed so that each point in one series is paired with one or more relevant points in the other series. Gait dissimilarities were calculated for both individual joint kinematics and the coordinated movement patterns of the lower limb. For the coordinated patterns, we calculated dissimilarities based on three-dimensional time series of joint kinematics (i.e., ankle, knee, and hip joint angles). Additionally, we calculated baseline dissimilarity as the mean dissimilarity across all possible pairs of the last five blocks of the baseline phase (i.e., _5_*C*_2_ = 10 pairs) for each participant, which was assumed to reflect the inherent variability of gait kinematics.

**Fig. 2.**
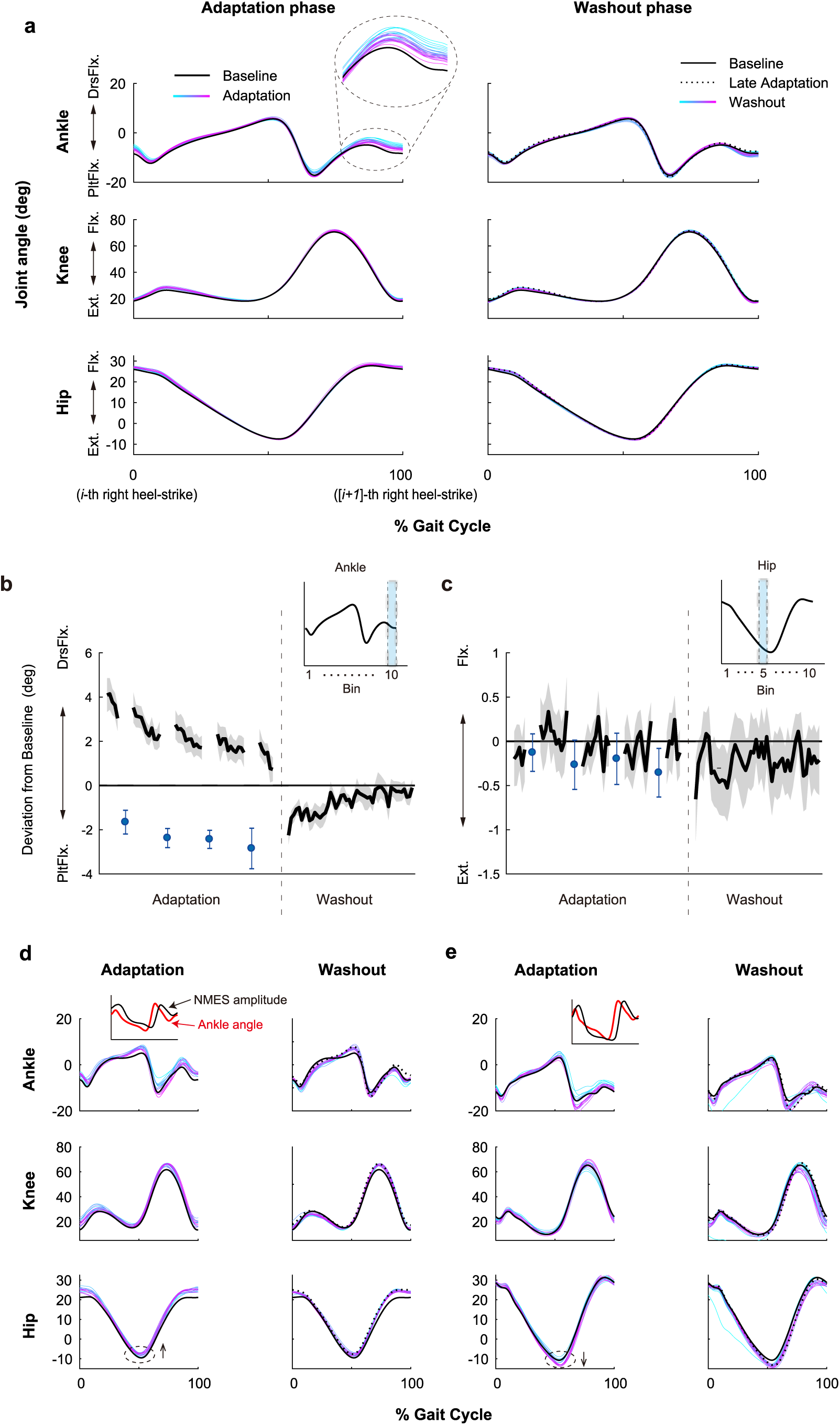
Changes in joint kinematics during the adaptation and washout phases. (a) Average changes in joint kinematics across participants over a gait cycle. DrsFlx: Dorsiflexion; PltFlx: Plantarflexion; Flx: Flexion; Ext: Extension. The line color changes from cyan to magenta as the blocks within each phase progress. The black solid and dashed lines represent the baseline and late adaptation kinematics, respectively. (b, c) Deviations from baseline in joint angles at specific time bins within the gait cycle. Each gait cycle was divided into 10 time bins, and the deviation within each bin was plotted. (b) The ankle joint angle around heel-strike (e.g., the 10th bin, shown in panel a) exhibited a clear tendency to maintain the baseline angle. (c) In contrast, other joints or time bins (e.g., the hip joint angle in the 5th bin, shown in panel c) did not display a consistent trend. Plots for all time bins of the three joints are shown in Supplementary Figure 1. The black line and shaded area represent the mean deviation from baseline and the standard error of the mean (SEM) across participants. Blue dots and error bars indicate the mean deviation and SEM during the catch blocks. (d, e) Individual joint kinematics for two representative participants, plotted in the same format as in (a). Black arrows indicate the direction of adaptive changes around peak hip extension (dashed circle). Insets in (d) and (e) show the average ankle joint angle (red) and NMES amplitude (black) during the last block of the adaptation phase for each participant. For clarity, the ankle joint angle trajectory in the insets is shown with its sign inverted relative to the main plot, so that increases in ankle angle (greater plantarflexion) correspond to increases in NMES amplitude.

**Fig. 3.**
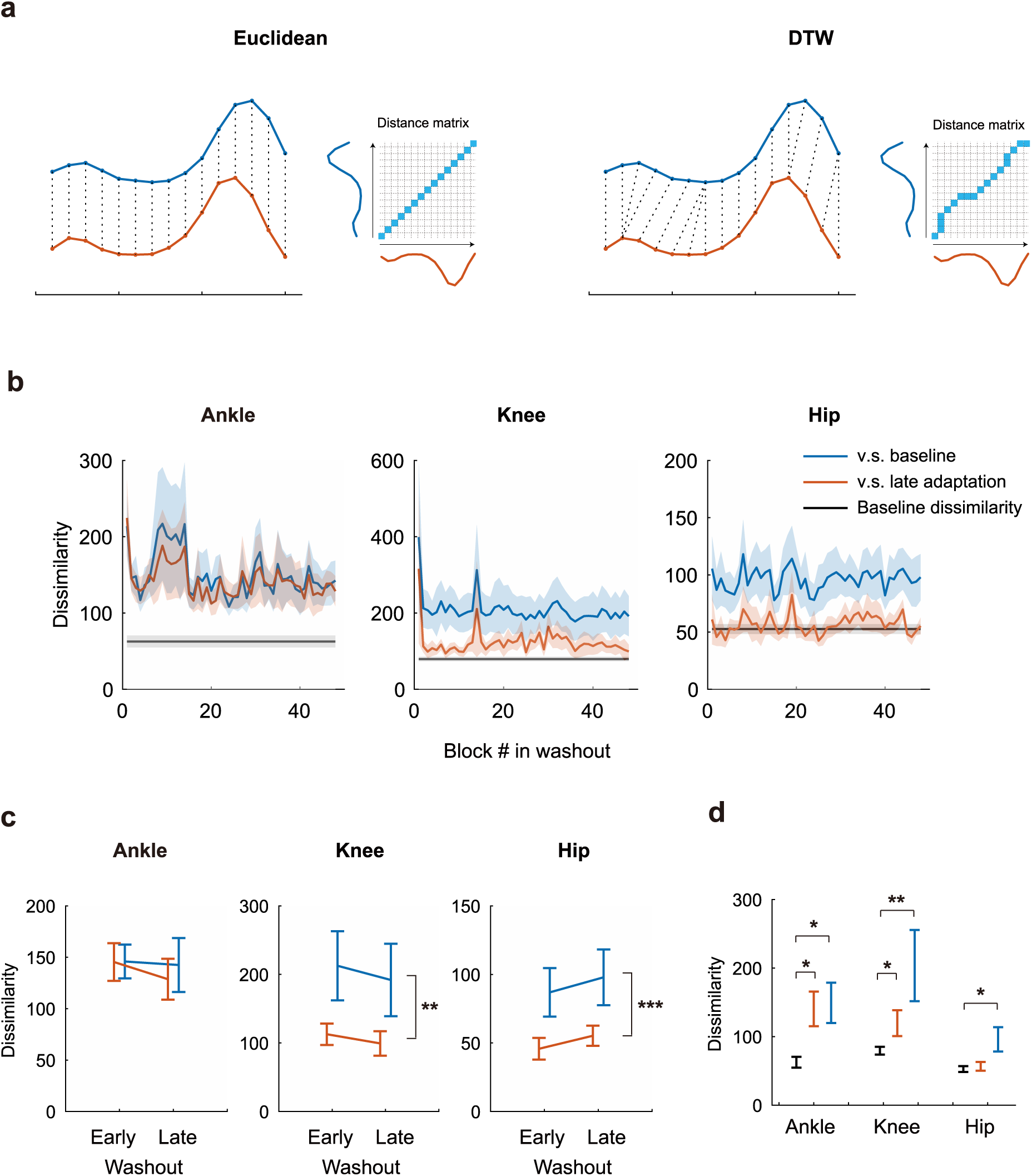
Gait dissimilarity analysis for single joints. Using the dynamic time warping algorithm (DTW), we calculated gait dissimilarities for each block of the washout phase relative to baseline and to late adaptation, as well as dissimilarities within baseline gait (baseline dissimilarities). (a) Schematic illustration comparing dissimilarity calculations based on Euclidean distance and the DTW algorithm. In the Euclidean distance-based method (left), dissimilarity is computed as the sum of point-to-point Euclidean distances between two time series (red and blue lines), where each point in one series is aligned with a single corresponding point in the other (dotted lines. In other words, the dissimilarity corresponds to the sum of the anti-diagonal elements of the distance matrix.) In contrast, DTW calculates the sum of distances between matched points in two series, where matching is performed to find the path through the distance matrix, from bottom-left to top-right, that minimizes the total cumulative distance (right). (b) Change in gait dissimilarity during the washout phase for each joint. Blue and red lines represent dissimilarities between washout and baseline, and between washout and late adaptation, respectively. Black lines represent the baseline dissimilarities. Shaded areas indicate the standard error of the mean (SEM) across participants. (c) Results of a two-way repeated-measures ANOVA on gait dissimilarity. Asterisks denote a significant main effect of phase (i.e., a difference between dissimilarities to baseline and dissimilarities to late adaptation) (**: p < 0.01; ***: p < 0.001). (d) Comparison of the baseline gait dissimilarities with those for washout vs. baseline and washout vs. late adaptation. Error bars indicate SEM across participants. Asterisks denotes significant differences (*: p < 0.025; **: p < 0.005).

### Statistical analysis

To examine the effects of phase (i.e., comparison to baseline or to late adaptation) and block (i.e., early or late block in the washout phase) on gait dissimilarity, we conducted a two-way repeated-measures ANOVA on gait dissimilarity. Prior to the analysis, we performed an aligned rank transformation on the dissimilarity values, which enables non-parametric comparison analogous to a factorial ANOVA [Wobbrock et al., 2011].

We defined the early and late blocks of the washout phase as the second and 48th blocks, respectively. The second block was selected as the early block because data from the first block showed high variability due to the abrupt removal of the perturbation. We initially planned to perform post hoc tests if an interaction effect was detected; however, no interaction effect was found. To compare the dissimilarity between washout and either baseline or late adaptation with the inherent variability, we conducted Wilcoxon signed-rank tests to detect potential differences. For this comparison, we used the average dissimilarities across the second through 48th blocks. Except for the comparison to inherent variability, the significance level alpha was set to 0.05. For the comparison to the inherent variability, the significance level alpha was adjusted to 0.025 using the Bonferroni correction to control for multiple comparisons. All values are reported as mean ± standard error. All statistical analyses were conducted using R version 4.4.2.

## Results

### Subject-averaged and individual gait pattern changes during adaptation and washout

Figure 2 illustrates how the gait pattern changed throughout the task. Subject-averaged joint kinematics indicate that the gait pattern changed in response to the NMES perturbation, primarily at the ankle joint, which was directly perturbed by NMES (Fig. 2a, left). Specifically, the initial excessive dorsiflexion induced by NMES gradually decreased during adaptation. In addition, greater flexion was observed in the knee and hip joints compared to baseline; however, these changes were smaller than those in the ankle joint. During the washout phase, kinematics generally returned to baseline levels, partly because they were already close to baseline at the start of the washout phase (Fig. 2a, right).

To investigate the joint kinematic changes in more detail, we divided the gait cycle averaged within each block of the adaptation and washout phases into ten time bins and analyzed changes in joint angles from baseline within each bin. For the ankle joint angle around heel-strike, we observed a typical adaptation–washout pattern commonly seen in planar reaching tasks: an initial perturbation-induced deviation that gradually diminished during adaptation, followed by a deviation in the opposite direction in the washout phase, which again decreased over time (Fig. 2b). These results suggest that, in response to excessive dorsiflexion induced by NMES, the motor system attempted to restore the ankle joint angle during heel-strike to its baseline level. In contrast, no clear patterns were observed in the other time bins or joints (Fig. 2c; joint angle changes for all time bins and joints are shown in Supplementary Fig. 1).

While consistent changes in the subject-averaged kinematics were observed only at specific time bins and joints, individual gait patterns exhibited more pronounced changes (Fig. 2d, e). The joint kinematics of two representative participants revealed broader, individual-specific adaptations not only at the ankle but also at other joints during the adaptation phase. For example, adaptive changes in the hip joint angle at peak extension (dashed circle in Fig. 2d, e) occurred in opposite directions in the two participants. Furthermore, and importantly, the gait patterns in the washout phase remained closer to those observed at the end of the adaptation phase (dashed line) rather than to those at baseline (solid line) for both participants, in contrast to the typical findings from standard planar reaching tasks.

These observations suggest that the possible changes in kinematics induced by NMES could be obscured in the averaged patterns and that they should be evaluated using metrics that reflect intra-individual changes in gait patterns rather than relying solely on subject-averaged kinematics (see the next section).

### Gait dissimilarity analysis reveals group-Level persistence of adapted gait patterns

To further investigate the characteristic behavior observed in the two representative participants (Fig. 2d, e), we assessed, for each participant, the extent to which joint kinematics in each block of the washout phase differed from those of baseline or late adaptation (i.e., dissimilarity). This approach allows us to characterize the pattern of changes in gait kinematics during the washout phase, irrespective of inter-individual differences.

We quantified gait pattern dissimilarity using the DTW algorithm (Fig. 3a). Overall, dissimilarities between washout and baseline were greater than those between washout and late adaptation, and this pattern remained consistent throughout the washout phase (Fig. 3b). This suggests that gait patterns during the washout phase were more similar to those observed during late adaptation than to those of baseline. This finding is consistent with the observations from the two representative participants, whose gait patterns did not return to baseline for at least 8 minutes and remained similar to those observed during late adaptation (Fig. 2d, e). Importantly, these trends, which were not apparent when simply averaging kinematics across participants, became evident at the group level when analyzed based on individual gait pattern dissimilarities.

In detail, the trends differed between the ankle and other joints (Fig. 3b). To statistically evaluate the similarities between washout and late adaptation gait patterns as well as their persistence, we performed a two-way repeated-measures ANOVA with aligned rank transformation on gait dissimilarity, with phase (vs. baseline or vs. late adaptation) and block (early [2nd] or late [48th] block in the washout phase) as main effects (Fig. 3c). For the ankle joint, there was no significant main effect of block (F[1, 27] = 0.721, *p* = 0.403) or phase (F[1, 27] = 0.037, *p* = 0.850), nor a significant interaction effect (F[1, 27] = 0.048, *p* = 0.828). However, for the knee and hip joints, a significant main effect was found for phase (F[1, 27] = 13.315, *p* = 0.00111 for the knee; F[1, 27] = 33.690, *p* = 3.549 * 10^-6^ for the hip), but not for block (F[1, 27] = 1.549, *p* = 0.224; F[1, 27] = 1.746, *p* = 0.197) or the interaction effect (F[1, 27] = 0.200, *p* = 0.658; F[1, 27] = 0.019, *p* = 0.890). The absence of a significant main effect of phase at the ankle joint is likely due to the direct application of the perturbation torque to the ankle: its removal inevitably altered ankle joint kinematics, causing the joint kinematics during the washout phase to differ not only from baseline but also from late adaptation.

Additionally, we calculated the average dissimilarities among baseline gait patterns for each participant, which are assumed to reflect individual inherent variability in gait. We then compared them to the dissimilarities between washout and either late adaptation or baseline (Fig. 3a, c). This comparison allows us to assess whether the observed dissimilarities between washout and baseline exceed the level of natural variability present during unperturbed walking. Multiple pairwise comparisons revealed that the average dissimilarities between washout and baseline were significantly greater than the inherent variability for all joints (*p* = 0.004, 0.002, and 0.010 for the ankle, knee, and hip, respectively). In contrast, the dissimilarities between washout and late adaptation were not significantly different from inherent variability for the hip joint (*p* = 0.922), but were significantly greater than inherent variability for the ankle and knee joints (*p* = 0.014 and 0.020, respectively).

Taken together, these analyses suggest that, at the group level, the gait pattern did not return to baseline after the perturbation was removed. Instead, it tended to retain kinematics of the late adaptation, particularly in the knee and hip joints, despite individual differences in the specific kinematic changes.

### Inter-joint coordination also did not return to baseline after perturbation removal

Next, we extended our analysis from individual joints to inter-joint coordination. To investigate coordination among the hip, knee, and ankle joints, we plotted the phase trajectories of the three joint angle changes over a gait cycle in three-dimensional space for both the group-averaged data (Fig. 4a) and a representative participant (Fig. 4b). As in the single-joint analysis, the representative data showed that the joint coordination pattern did not return to baseline after perturbation removal, although this tendency was less pronounced when averaged across participants.

**Fig. 4.**
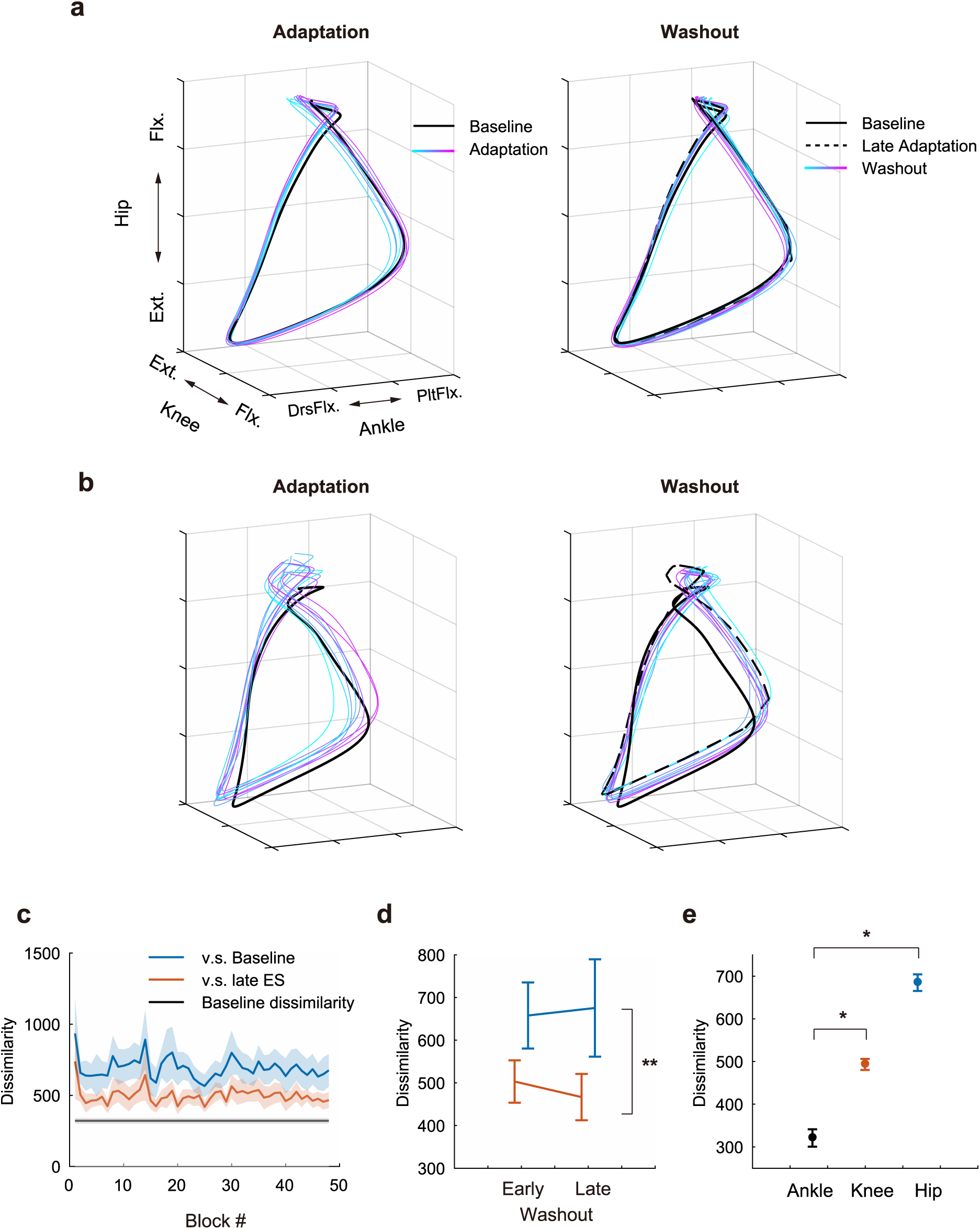
Analysis of inter-joint coordination in gait. (a, b) Changes in joint coordination during the adaptation and washout phases. Trajectories of three joint angles in one gait cycle are plotted. Averaged data across participants (a) and data from the representative participant (b) are shown. As in Figure 2, the line color changes from cyan to magenta as the blocks within each phase progress. The black solid and dashed lines represent the baseline and late adaptation kinematics, respectively. (c–e) Dissimilarity analysis of coordinated gait patterns, based on the three joint angles over one gait cycle. (c) Change in gait dissimilarity during the washout phase. Blue and red lines represent dissimilarities between washout and baseline, and between washout and late adaptation, respectively. Black lines represent the baseline dissimilarities. Shaded areas indicate the standard error of the mean (SEM) across participants. (d) Results of a two-way repeated-measures ANOVA on gait dissimilarity. Asterisk indicates a significant main effect of phase (**: p < 0.01). (e) Comparison of the baseline gait dissimilarities with those between washout and either baseline or late adaptation. Error bars indicate SEM across participants. Asterisks indicate significant differences (*: p < 0.025).

To quantify the extent to which coordinated gait patterns in the washout phase resembled those at baseline or late adaptation for each individual, we applied the same dissimilarity-based approach used for single-joint kinematics (Fig. 4c). Dissimilarity analysis revealed that the overall gait pattern in the washout phase was more similar to that of late adaptation, with a significant main effect of phase (F[1, 27] = 8.168, *p* = 0.008), and showed no systematic change over time (no significant main effect of block or interaction effect; F[1, 27] = 0.1411, *p* = 0.710; F[1, 27] = 0.0415, *p* = 0.840). When compared with inherent variability, dissimilarities were significantly greater for both washout vs. late adaptation (*p* = 0.014) and washout vs. baseline (*p* = 0.004) (Fig. 4e). Thus, the coordinated gait pattern in the washout phase also tended to retain features of late adaptation kinematics.

## Discussion

In this study, we investigated adaptive changes in gait kinematics in response to NMES-based perturbation on the ankle dorsiflexor muscle (TA). The present results indicate that gait adaptation to NMES perturbation involved a global modification of the walking pattern, not limited to the directly perturbed ankle joint. Furthermore, these adaptive changes in gait kinematics tended to persist even after the perturbation was removed, rather than returning to the baseline pattern.

An important aspect of this study is its focus on changes in joint kinematics over the entire washout period. Many gait adaptation studies, including those employing split-belt treadmill adaptation paradigms, have evaluated adaptation with summary measures such as gait asymmetry or energy expenditure, rather than directly analyzing joint kinematics. However, such measures are insufficient to capture how the motor system coordinates redundant degrees of freedom to achieve task goals, as identical summary values do not necessarily correspond to identical joint-level gait patterns. Several studies have reported prolonged aftereffects in gait adaptation [Selinger et al., 2015; Luu et al., 2017], suggesting that gait may not fully return to baseline after perturbation removal. However, as these were evaluated using summary measures, how joint kinematics change during washout under kinematic redundancy remains unclear. Conversely, gait adaptation studies examining joint kinematics often lack detailed analysis of the washout phase, with available data limited to its early portion or a few discrete time points [Lam et al., 2006; Kambic et al., 2023; Kim et al., 2010], thereby hindering a comprehensive understanding of how joint kinematics change throughout the washout phase. By analyzing the temporal evolution of joint kinematics across the entire washout period, the present study enabled a more detailed investigation of how the motor system recalibrates its motor plan under kinematic redundancy after perturbation removal.

### Mechanisms underlying the persistence of adapted gait patterns

Our finding that gait kinematics did not necessarily return to baseline following perturbation removal can be attributed to task redundancy. The nature of a redundant system, where the same task goal can be achieved by multiple joint kinematics, allows the motor system to select a movement pattern different from baseline during the washout phase. However, even in a redundant system, a return to baseline would still be possible, suggesting that a gait pattern different from the baseline was selected for some reason.

As one possible explanation for this, we propose a hypothetical framework based on the concept of optimal control (Fig. 5). A body of prior studies suggests that the motor system selects an ‘optimal’ movement that minimizes a certain cost from an infinite number of possible choices for accomplishing a given task [Flash and Hogan, 1985; Harris and Wolpert, 1998; Todorov and Jordan, 2002; Todorov, 2004; Izawa et al., 2008; Emken et al., 2007; Finley et al., 2013]. Looking at the current task situation from this perspective, the gait pattern at baseline can be regarded as an optimal, or at least near-optimal, solution based on the cost landscape determined by factors such as individual musculoskeletal properties and walking speed (Fig. 5a). Once a perturbation is applied, the cost function landscape is altered, and the motor system acquires a new optimal kinematic pattern based on the new cost landscape (Fig. 5b). When the perturbation is removed, the cost function landscape returns to its original form at baseline, and the motor system recalibrates its motor plan within this restored landscape (Fig. 5c). During this process, if the motor system explores movement solutions over a sufficiently wide range, the gait pattern returns to baseline, which is optimal under the original cost landscape (Fig. 5c, top). Conversely, if the exploration range is limited to a neighborhood of the kinematics acquired under NMES, the resulting gait pattern may become trapped in a local optimum, thus resembling the adapted pattern (Fig. 5c, bottom).

**Fig. 5.**
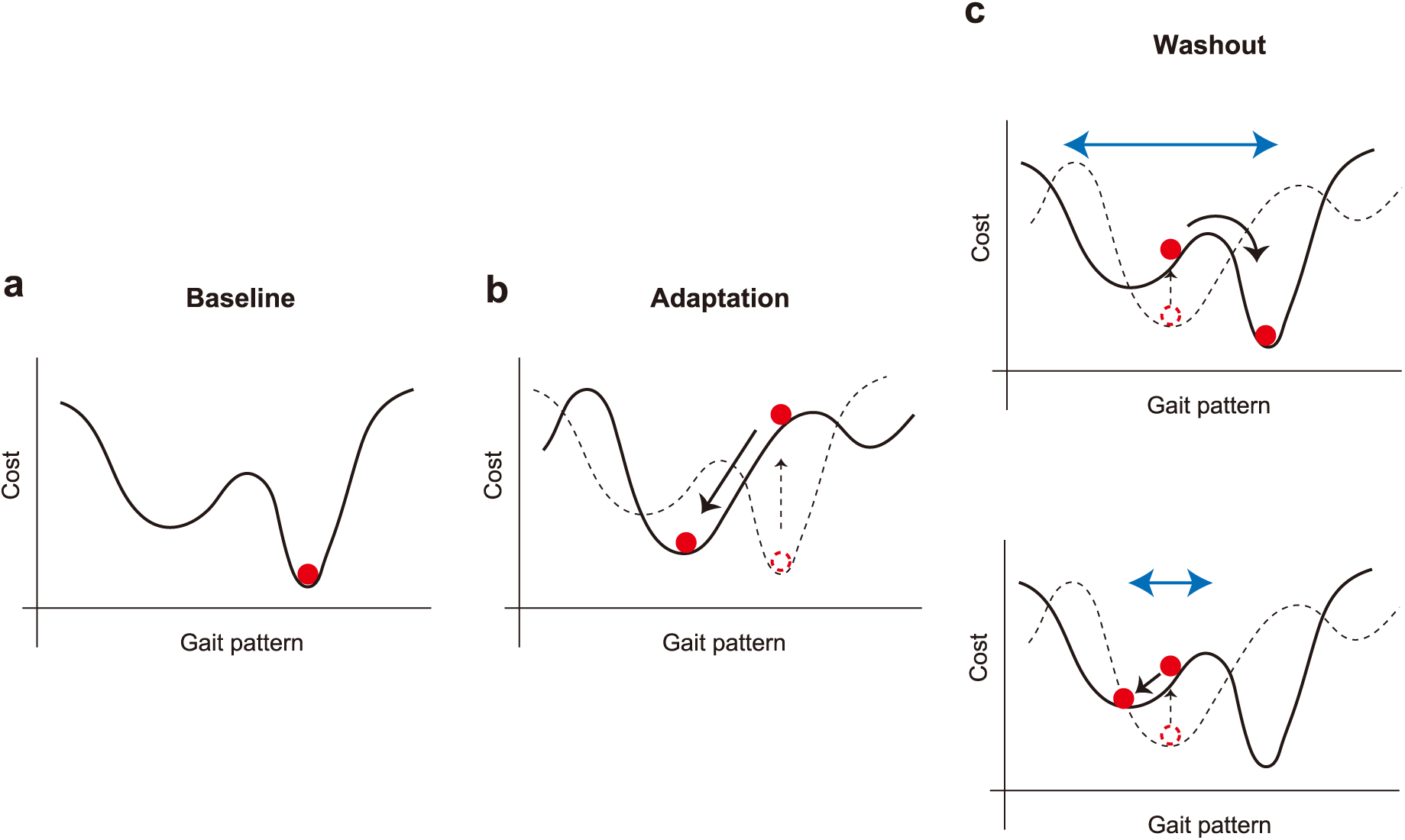
Schematic illustration of the framework based on optimal control theory. (a) In the baseline phase, the gait pattern is considered the optimal solution under the cost landscape specific to an individual (red circle). (b) During the adaptation phase, the manipulation of musculoskeletal dynamics by the perturbation alters the cost landscape (dashed line to solid line), and the gait pattern is re-optimized based on the new landscape (black arrow). (c) In the washout phase, the cost landscape returns to its original form (dashed line to solid line), and the gait pattern is recalibrated within the original landscape. (top) If the motor system explores the landscape sufficiently broadly (blue arrow), the recalibrated gait pattern should be identical to the baseline pattern. (bottom) In contrast, a limited range of exploration (blue arrow) may result in different gait patterns after the removal of the perturbation. The shape of the cost landscape would also influence whether the gait pattern returns to baseline.

Although this framework is speculative and it remains unclear how the brain explores movement solution space to optimize performance, it aligns with the experimental results observed in this study. To investigate the validity of this framework, predictive simulation of gait using a musculoskeletal model will be a promising approach. By incorporating NMES perturbation into a musculoskeletal model and systematically manipulating the range of exploration during predictive simulation, it will be possible to examine the relationship between the extent of exploration and the resulting gait patterns.

The persistence of NMES-induced kinematics during the washout phase intuitively suggests the possible involvement of use-dependent plasticity (UDP) [Classen et al., 1998; Diedrichsen et al., 2010; Wood et al., 2020], where the repetition of a specific movement induces plastic changes in the activity of cortical motor areas, thereby biasing subsequent movements toward the repeated movement. From this perspective, the repetition of the gait pattern acquired under NMES may have induced UDP, which in turn contributed to the persistence of a similar gait pattern in the washout phase. However, given that UDP likely induces plastic changes at the level of muscle activation rather than kinematics, it is unlikely to be the sole mechanism underlying the persistence of NMES-induced kinematics: maintaining the muscle activation acquired under NMES after its removal would likely have resulted in different kinematics from the NMES-induced pattern.

Additionally, this persistent change in gait pattern may be related to NMES-induced changes in corticospinal excitability, as suggested by previous literature [Ridding et al., 2000; Khaslavskaia et al., 2002; Nuara et al., 2023]. To determine whether this is a specific effect of NMES or a general property of redundant systems, studies using a similar experimental paradigm with alternative perturbation methods, such as mechanical perturbation with an exoskeleton, would be valuable.

### Gait pattern changes in adaptation to NMES perturbation

There appears to be no consistent pattern of adaptive changes in the overall gait pattern across participants. However, when focusing on specific joint movements or phases of the gait cycle, several consistent tendencies emerged. In particular, we observed a consistent tendency to maintain the ankle joint angle around heel-strike although NMES intensity was set to be maximal at peak plantarflexion, which likely occurs shortly after toe-off. This temporal gap of several hundred milliseconds can be partly attributed to the electromechanical delay (EMD) [Cavanagh and Komi, 1979], as well as latency inherent in the perturbation system. EMD, the delay between the onset of muscle activation (in this case, the onset of NMES) and the actual production of torque, likely caused the peak perturbation torque to occur closer to heel-strike. In addition, deviation from the appropriate ankle joint angle at heel-strike can disrupt smooth forward progression of the center of mass. Hyper-dorsiflexion at heel-strike increases heel rocker function, which leads to excessive forward rotation of the tibia over the foot, thereby disrupting a controlled loading response [Perry and Burnfield, 2010]. On the other hand, excessive plantarflexion reduces the heel rocker function [Attias et al., 2023], thereby diminishing shock absorption during initial contact. These biomechanical constraints may have contributed to the observed tendency to maintain the baseline ankle angle at heel-strike. Taken together with the diverse patterns of overall gait adaptation observed across individuals, this suggests that the motor system may attempt to achieve both local and global optimization simultaneously.

### Potential of NMES as a method for movement modification

The persistence of the adapted gait pattern in the washout phase suggests that NMES perturbation has the potential to serve as a method to modify movement patterns in a given motor task. The present method modified already acquired movement patterns in healthy individuals within a short timescale, which contrasts with functional electrical stimulation (FES), a technique typically applied over longer timescales to restore impaired motor function. An advantage of this approach is its ability to implicitly alter movements. Due to the complexity of the multi-joint musculoskeletal system and the resulting interaction torques, intentionally altering a movement pattern can be challenging and may lead to unintended compensatory changes in other joints. Therefore, this implicit nature of NMES-based perturbation may be particularly beneficial for complex, real-world motor tasks.

To test the practical applicability of this method, future studies should examine the retention and generalization of its effects, including how long they persist and whether they remain stable across different contexts (e.g., transition from treadmill to ground walking). In addition, it is crucial to develop methods for predicting how movement patterns change in response to a given perturbation, taking inter-individual variability into account. These directions will be essential not only for practical applications but also for understanding how the motor system selects and calibrates movements from an infinite set during the processes of motor adaptation and de-adaptation.

### Differences between gait and goal-directed movements

It remains unclear whether the persistence of adaptive changes would also occur in other types of redundant goal-directed movements because these movements differ from gait in several fundamental ways. First, the kinematics of gait and goal-directed movements reflect different hierarchical structures of redundancy. In the present study, we focused on redundancy observed in joint kinematics. In goal-directed movements, this redundancy can generally be divided into two levels: trajectory-level redundancy (i.e., the existence of multiple end-effector trajectories for a given task) and joint-level redundancy (i.e., the existence of multiple joint movements for a given trajectory). In contrast, dissociating these two levels of redundancy in gait is challenging because it is difficult to define the equivalent of an end-effector trajectory that is present in goal-directed movements. Therefore, it is not possible to determine whether the persistent kinematic changes observed in the present study can be attributed to trajectory-level redundancy, joint-level redundancy, or both.

In addition, gait is a periodic and largely automatic behavior, with substantial contributions from central pattern generators in the spinal cord [Kiehn, 2006; Grillner, 2006]. Although supraspinal structures such as the cerebral cortex, cerebellum, and basal ganglia are also involved in gait control and adaptation [Takakusaki, 2013], the neural mechanisms underlying gait adaptation under redundancy may differ from those involved in discrete, voluntary movements like reaching.

Given these differences, future studies on joint kinematic responses to perturbations in redundant goal-directed tasks would be valuable for advancing our understanding of motor control and adaptation under kinematic redundancy. Recent research has developed redundant reaching tasks by adding extra degrees of freedom to planar reaching paradigms [Singh et al., 2016; Kobayashi and Nozaki, 2024]. These approaches offer a promising direction for comparing adaptation processes under kinematic redundancy across distinct motor tasks.

## Conclusion

In summary, we revealed that gait pattern changes resulting from adaptation to NMES perturbation tended to persist even after the perturbation was removed. These changes were not confined to the ankle joint but were observed globally across all three joints, despite the perturbation being applied specifically to that joint. These findings were demonstrated by examining kinematic changes during the washout phase, an aspect that has been relatively unexplored in prior studies. Although a detailed investigation of the mechanisms underlying this persistence is beyond the scope of the present study, our proposed optimal control-based framework provides a plausible explanation. These results not only highlight the unique characteristics of movement adaptation specific to redundant systems but also demonstrate that NMES-based perturbation is a promising approach for implicitly inducing changes in movement patterns.

## Acknowledgement

This work was supported by KAKENHI (21H04860 and 20H05459) and MTG Co., Ltd to D.N.

## Author contributions

Conceptualization, R.M. and D.N.; methodology, R.M. and D.N.; formal analysis, software, and investigation, R.M.; visualization, R.M. and D.N.; resources, D.N.; writing—original draft, R.M. ; writing—review and editing, R.M. and D.N.; funding acquisition, D.N.; supervision, D.N. All authors have read and agreed to the published version of the article.

---

**Supplementary Fig. 1.**
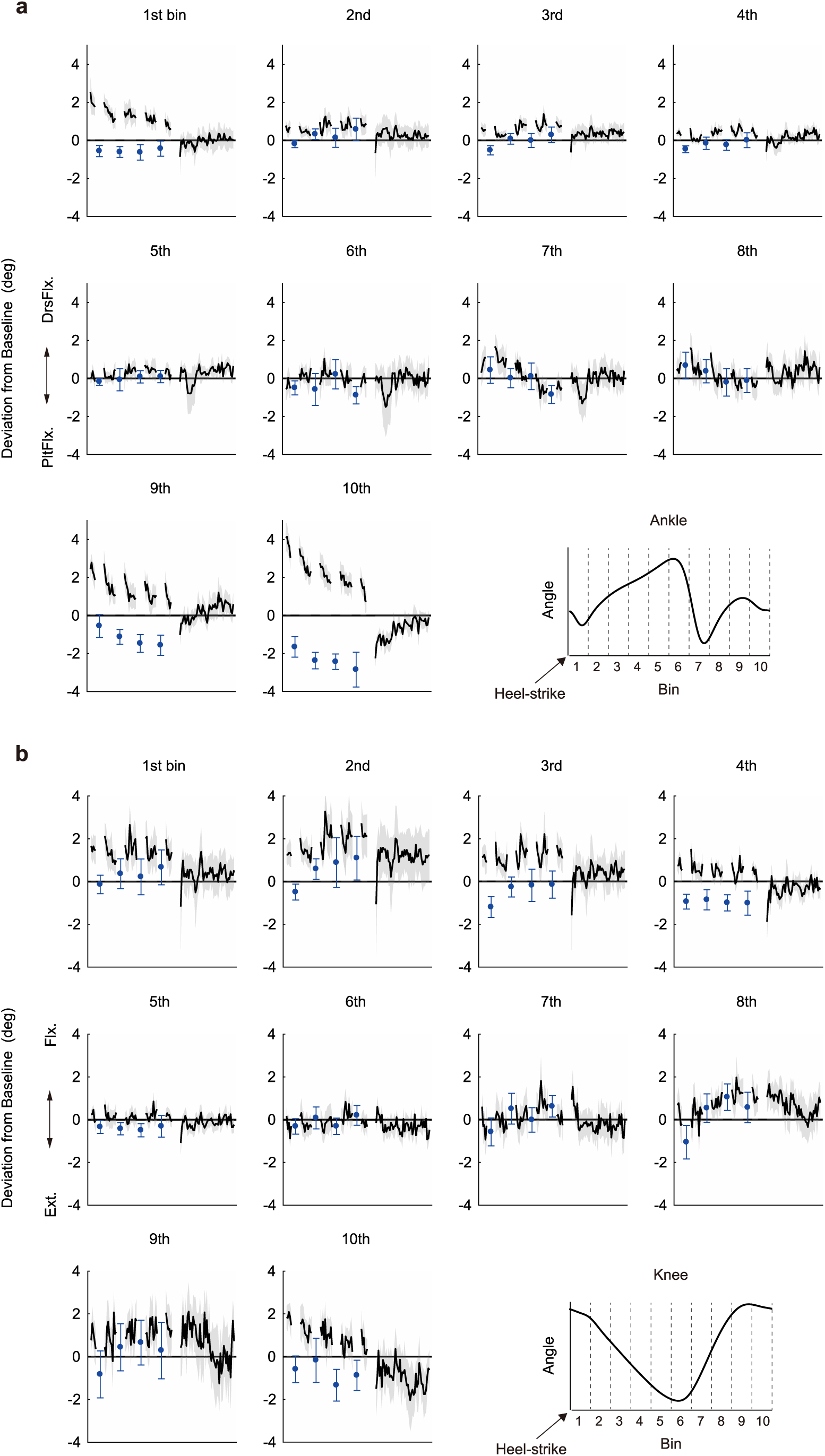
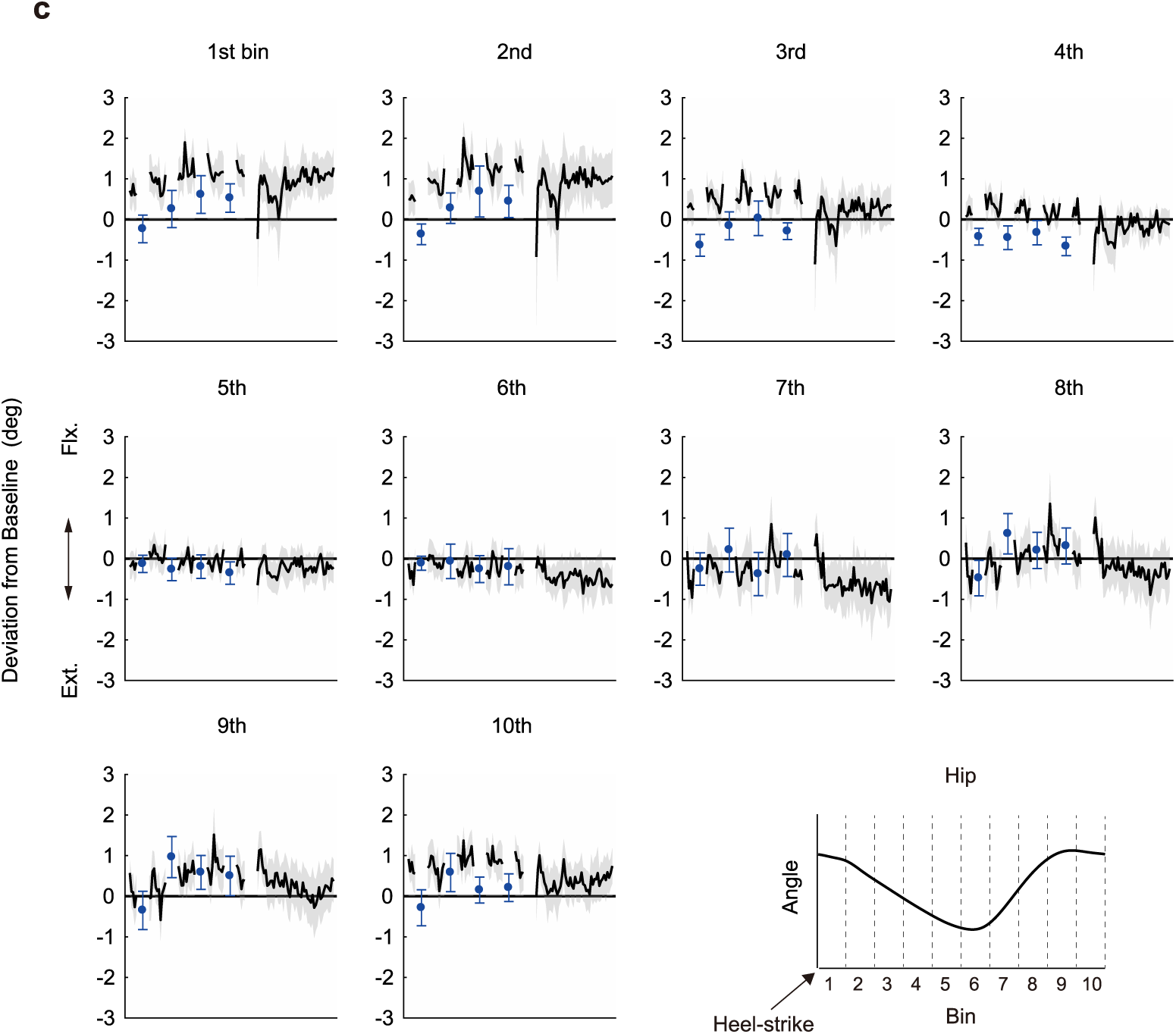
Deviations from baseline for the (a) ankle, (b) knee, and (c) hip joint angles at specific time bins of a gait cycle. The numbers above each plot correspond to the time-bin index. The 1st bin corresponds to the time point immediately after heel-strike. The black line and shaded area represent the average deviation from baseline and the standard error of the mean (SEM) across participants. Blue dots and error bars indicate the average deviation and SEM during the catch blocks.

